# Neural Dynamics of Executive Function in Cognitively-able Kindergarteners with Autism Spectrum Disorders (ASD) as Predictors of Concurrent Academic Achievement

**DOI:** 10.1101/455485

**Authors:** So Hyun Kim, George Buzzell, Susan Faja, Yeo Bi Choi, Hannah Thomas, Natalie Hiromi Brito, Lauren C. Shuffrey, William P. Fifer, Frederick D. Morrison, Catherine Lord, Nathan Fox

## Abstract

Although electrophysiological (EEG) measures of executive functions (EF) (e.g. error monitoring) have been used to predict academic achievement in typically developing (TD) children, work investigating a link between error monitoring, and academic skills in children with autism spectrum disorder (ASD) is limited. In the current study, we employed traditional electrophysiological and advanced time-frequency methods, combined with principal components analyses, to extract neural activity related to error monitoring, and tested their relations to academic achievement in cognitively-able kindergarteners with ASD. Thirty-five cognitively-able kindergarteners with ASD completed academic assessments and the child-friendly “Zoo Game” Go/No-go task at school entry. The Go/No-go task successfully elicited an error-related negativity (ERN) and error positivity (Pe) in children with ASD as young as 5 years at medio-frontal and posterior electrode sites, respectively. We also observed increased response-related theta power during errors relative to correct trials at medio-frontal sites. Both larger Pe and theta power significantly predicted concurrent academic achievement after controlling for behavioral performance on the Zoo Game and IQ. These results suggest that the use of time frequency EEG analyses, combined with traditional ERP measures, may provide new opportunities to investigate neurobiological mechanisms of EF and academic achievement in young children with ASD.

---

Recent estimates indicate that 512,000 students in the U.S. received educational services for autism spectrum disorder (ASD) under the Individuals with Disabilities Education Act in 2013 (Kena et al., 2015). With increasing early awareness and intervention efforts, greater numbers of children with ASD make significant cognitive and behavioral gains during the toddler and preschool years (e.g., Dawson et al., 2010) and are able to attend kindergarten with typically developing (TD) peers. Yet, we have limited knowledge about factors that contribute to academic outcomes in cognitively-able children with ASD. Prior work has employed behavioral assessment of executive function (EF; Fuster, 1997; Miyake et al., 2000) as one of the key predictors of academic outcomes of early literacy and math skills in TD children (Blair, 2002, 2016; Cameron et al., 2012; McClelland, Acock, & Morrison, 2006; McClelland et al., 2007; Ponitz, McClelland, Matthews, & Morrison, 2009; Willoughby, Kupersmidt, & Voegler-Lee, 2012). Children with ASD, compared to TD children, have significantly more difficulties with EF (Happé, Booth, Charlton, & Hughes, 2006; Konstantareas & Stewart, 2006), which may have cascading effects on their academic outcomes. Particular EF skills, such as “error monitoring,” can be assessed through the use of non-invasive neural measures, such as electroencephalography (EEG; Taylor, Stern, & Gehring, 2007). EF is traditionally divided into working memory, inhibitory control, and attention shifting; however, error monitoring has also been found to be an integral part of EF (Carter et al., 2000; Kramer et al., 2014), and neural activities related to error monitoring have been found to be critical for the development of academic functioning in the past (Hirsh & Inzlicht, 2010; M. Kim et al., 2016). Yet, we have limited knowledge about how error monitoring contributes to academic outcomes in cognitively-able children with ASD.

Error monitoring is not easily assessed via behavioral metrics; however, EEG-based markers of error monitoring are well established (Cavanagh & Frank, 2014; Ullsperger, Fischer, Nigbur, & Endrass, 2014), based on analysis of event-related potentials (ERPs; Luck & Kappenman, 2011) and time-frequency decompositions of the EEG signal (Cohen, 2014). When errors are made, ERPs derived from the EEG recording reveal more negative amplitudes over frontocentral scalp locations within ~100 ms of making an error, referred to as the “error-related negativity” (ERN; Falkenstein, Hohnsbein, Hoormann, & Blanke, 1991; Gehring, Goss, Coles, Meyer, & Donchin, 1993), along with a later emerging positivity (200-500 ms after making an error) over centroparietal scalp regions, known as the “error positivity” (Pe; Overbeek, Nieuwenhuis, & Ridderinkhof, 2005). The ERN reflects a rapid, automatic and likely unconscious aspect of error monitoring (Frank, Woroch, & Curran, 2005; Larson, Clayson, & Clawson, 2014; Luck & Kappenman, 2011), with the Pe reflecting a slower and more deliberative form of error monitoring that relates to decisions about what to do after an error (Gehring et al., 1993; Hajcak, McDonald, & Simons, 2003; Overbeek et al., 2005; Schneider, 2010; Steinhauser & Yeung, 2010). The ERN and Pe have been shown to at least partially involve anterior cingulate cortex (ACC) function although much more distributed activity is also present (Agam et al., 2011; Buzzell et al., 2017). The ACC has been implicated in error monitoring, as well as cognitive control functions more generally, such as detecting response conflict and attentional control, enabling the brain to adapt behavior to changing task demands and environmental circumstances (Botvinick, Cohen, & Carter, 2004; Posner & Raichle, 1994).

Two previous studies of a large sample of TD preschoolers successfully identified the presence of both an ERN and Pe using a child-friendly Go/No-Go task (Grammer, Carrasco, Gehring, & Morrison, 2014) and a significant association between these ERPs and academic achievement (M. Kim et al., 2016). In the study by M. Kim and colleagues (2016), in TD preschoolers as young as 3-5 years, increased Pe amplitudes predicted math achievement, but no association between the ERN and academic skills was identified. In contrast, other work in adults has found *both* the ERN *and* Pe to be significant predictors of academic achievement (Hirsh & Inzlicht, 2010). Extending this work to children with ASD, the ERN and Pe have been also been detected in children with ASD by our group based on a preliminary sample using the same child-friendly Go/No-Go task (S.H. Kim, Grammer, Benrey, Morrison, & Lord, 2017); similarly, other child-friendly tasks have been used to identify the presence of an ERN and Pe in children with ASD (Groen et al., 2008; Henderson et al., 2006).

Building directly on prior work establishing the presence of response-locked ERPs reflective of error monitoring (i.e., ERN and Pe) in both TD and children with ASD, the current study seeks to explore how error monitoring relates to academic achievement in kindergarteners with ASD. Although other ERP components might also relate to academic achievement in children with ASD, studies focusing on other ERP components such as N2 and P3 in kindergarten and preschool ages have resulted in mixed findings, with a handful of studies showing that N2 and/or P3 can be elicited in preschool children (e.g., Abdul Rahman, Carroll, Espy, & Wiebe, 2017; Grabell, Olson, Tardif, Thompson, & Gehring, 2017), while others found that P3 might not be present in children younger than 8-10 years of age, and N2 might be widely distributed across the frontal and parietal electrodes (Buss, Dennis, Brooker, & Sippel, 2011; Jonkman, 2006). Therefore, given prior work that has specifically validated the study of ERPs associated with error-monitoring in young children and their associations with academic outcomes (Grammer et al., 2014; M. Kim et al., 2016; S. H. Kim et al., 2017), the hypotheses of the current studies are similarly focused on the ERN and Pe. Specifically, we hypothesized that an enhanced Pe observed in the posterior sites, but not ERN in the frontal sites, would predict better academic achievement in cognitively more-able kindergarteners with ASD.

As mentioned above, a complementary approach to ERP assessments of error monitoring involves time-frequency (TF) decompositions of the EEG signal (i.e. conversion of raw EEG data into a representation of how neural oscillations change over time) to quantify how power within particular frequency bands (i.e. the strength of neural oscillations) changes following errors (Cohen, 2014). Theta oscillations (~4-8 Hz) are thought to reflect activation of a neural system responsible for monitoring and adapting behavior in response to events like errors (Cavanagh & Frank, 2014). Work in adults has shown such theta oscillations can be recorded using EEG electrodes located over the medio-frontal cortex, and found that theta power (strength of these oscillations) increases in response to error commission (Cavanagh & Frank, 2014). In the present study, we employed the TF approach in addition to analyses of traditional ERPs, because the TF approach may provide a more robust measurement of neural activity associated with error monitoring (and performance monitoring more generally). Furthermore, TF measures are less susceptible to artifact and noise, providing additional advantages for analyses of data obtained from children (Bowers et al., 2018). Thus, analyzing time-frequency measures of error monitoring can also provide additional, band-specific information, which might afford improved statistical power in predicting academic achievement in children with ASD over approaches that only analyze ERPs. For example, it has recently been shown that age-related changes in theta power associated with performance monitoring could be identified in the absence of similar changes in associated ERPs (Bowers et al., 2018). However, while studying error-related theta band dynamics is well established in adults (e.g., Cavanagh & Frank, 2014; Harper, Malone, & Bernat, 2014), much less work has been conducted in children (but see DuPuis et al., 2015 for a recent example). To our knowledge, work linking time-frequency measures to academic skills in children with ASD is non-existent. Given the previous research with adults and adolescents, we expected that larger theta power would be associated with better behavioral performance on EF tasks and academic skills in children with ASD.

Past studies indicate that EF-related neural measures using electrophysiological methods, specifically measures of error monitoring, could predict academic achievement in TD children and adults (Hirsh & Inzlicht, 2010; M. Kim et al., 2016). However, such links have not been studied in cognitively-able children with ASD. The examination of how error monitoring is associated with concurrent academic achievement in young children with ASD has important implications for educational planning and treatment programming aimed at improving early school success and subsequent longer term outcomes in this population. Therefore, the current study employed a series of neural measures, including the ERN and Pe ERPs, as well as time-frequency analyses of theta power, in an effort to better characterize the effects of error monitoring on academic skills in children with ASD.

## Method

### Participants

Participants included 35 cognitively-able 4 to 5-year-old children with ASD (Mean age=62.3 months, SD=4.4 months; 25 males) without general cognitive delays (full-scale IQ≥85; Mean IQ=104.7, SD=14.2) and without moderate to severe structural language delays (i.e., child must be using full, complex sentences). These children were assessed at kindergarten entry (Summer/Fall of 2017). A previous diagnosis of ASD was required as inclusion criteria; and the diagnosis of ASD was confirmed with the gold standard diagnostic measure, the Autism Diagnostic Observation Schedule-2 (ADOS-2; Lord et al., 2012). The ADOS-2 was carried out by examiners who achieved research reliability and were under the supervision of a licensed clinical psychologist. Comparison Scores on the ADOS diagnostic algorithms ranged from 3 to 10 (M=7.71, SD=1.9; Gotham, Pickles, & Lord, 2009). The sample included 16 (46%) white, 4 (11%) African American, 6 (17%) bi-racial, and 2 (6%) Asian children. The rest of the children were reported to be of other races (n=7, 20%). About 85% of caregivers had a bachelor’s degree or above (n= 30). Out of 35 children, 10 (29%) children were enrolled in general education classrooms; 7 (20%) were in inclusion classrooms (children with ASD are taught together with TD children); and 18 (51%) were in special education classrooms in schools in urban and suburban areas of New York. Based on parent report, twenty-eight (80.0%) children were right handed, 5 left (14.3%) and the rest were ambidextrous (n=2, 5.7%). No participants were taking medication at the time of testing.

### Electrophysiological Tasks and Measures

#### EEG/ERP Task

ERP/EEG patterns related to error monitoring (ERN, Pe, theta power) were measured based on a child-friendly Go/No-go task (“Zoo Game”; Grammer et al., 2014; Lamm et al., 2014) in a testing room with minimal distractions. Children were told that they are playing a game to help a zookeeper catch all the loose animals in the zoo except for three friendly orangutans who are helping the zookeeper. Children were asked to press a button as quickly as possible when they saw an animal (Go trials) but inhibit their responses when they saw an orangutan (No-go trials). A child started the game with a practice block of 12 trials (9 animals; 3 orangutans) followed by 8 blocks of the task, each with 40 trials, for a total of 320 trials (240 Go and 80 No-go trials). Each image was preceded by a fixation cross displayed for a randomized interval ranging from 200-300 ms. The stimuli were presented for 750 ms, followed by a blank screen for 500 ms. Responses could be made while the stimulus was on the screen or at any point during the following 500 ms (Figure 1). This task successfully elicited specific ERP components of interest (e.g., ERN, Pe) in a large sample of TD children as young as 3 years (Grammer et al., 2014). Each block consisted of novel sets of animal photographs, and each set is balanced with respect to color, animal type, and size. Children were given performance feedback of either “Try to catch them even faster next time,” if no-go trial accuracy was greater or equal to 90%, or “Watch out for the orangutan friends,” if no-go trials accuracy was less than 90%, after each block of the task (not by trial). These prompts were given to the children based on the calculation of the error rates to ensure an adequate number of trials for stable ERP/EEG waveforms. Children were allowed to have “Wiggle Time” between blocks. From the Zoo Game, we extracted the number and percentage of error/correct trials and reaction times.

**Figure 1.**
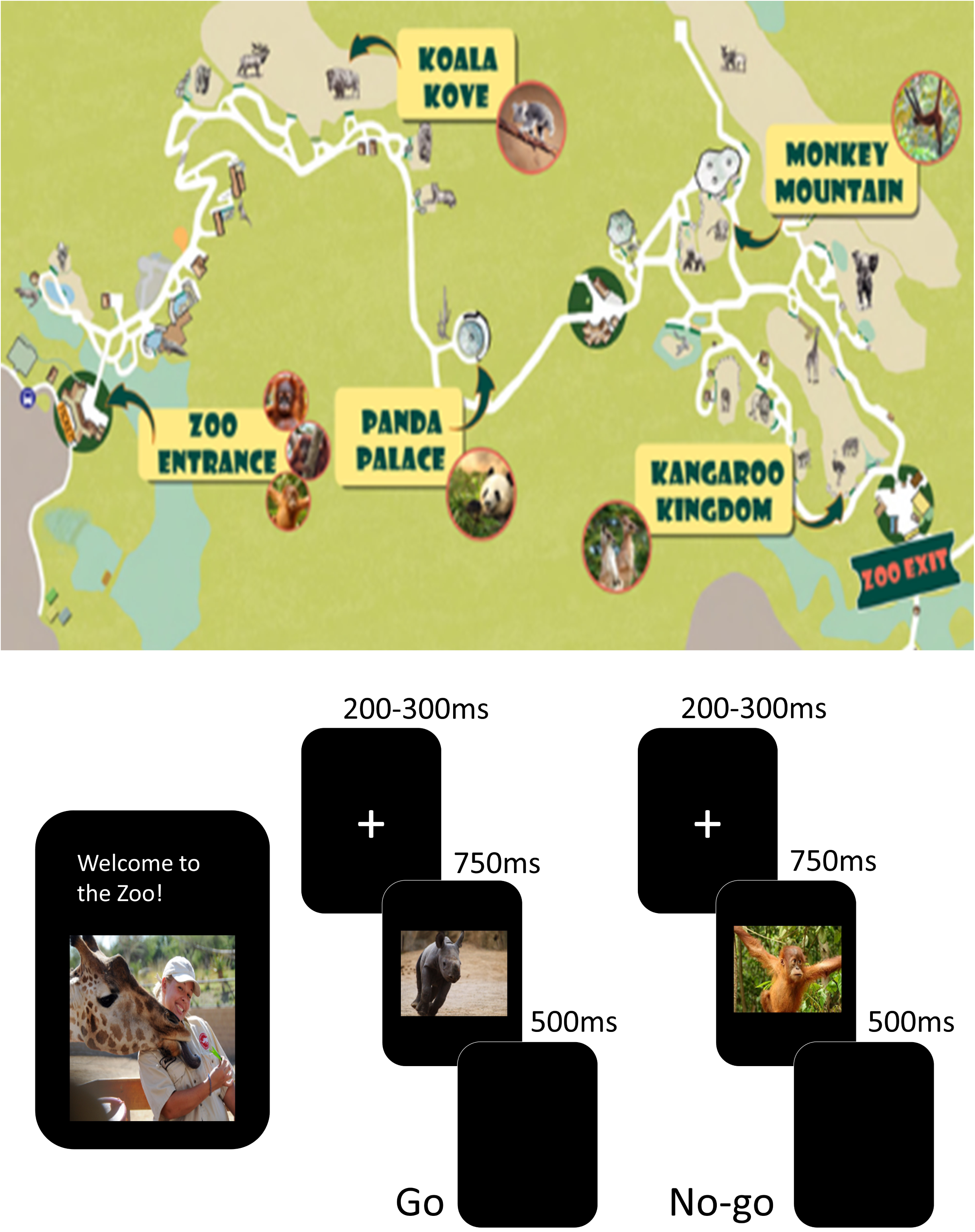
The Go/No-Go Zoo Game (Grammer et al., 2014). Children were shown the Zoo map at the beginning and in between blocks to show them the progress they were making on the game (upper half). The bottom half shows the design of the task including Go trials on the left (without an orangutan) and No-Go trials on the right (with an orangutan).

### Electrophysiological recording, data reduction, and data processing

Stimuli were presented on a PC laptop using E-Prime 2.0 software. Displays were viewed in a testing room with minimal distractions. EEG was recorded using Net Station 5.4 on a Macintosh laptop. A 64-channel HydroCel Geodesic sensor net (EGI) was soaked with potassium-chloride electrolyte solution, placed on the participant’s head, and fitted according to the manufacturer’s specifications. As recommended for this system, impedances were kept below 50 kΩ. The EEG signal was digitized and sampled at 500 Hz via a preamplifier system (EGI Geodesic NA 400 System). Data was processed offline using Matlab (The Mathwoks, Natick, MA), the EEGLAB toolbox (Delorme & Makeig, 2004) and custom Matlab scripts partly based on work by Bernat and colleagues (Bernat, Williams, & Gehring, 2005). EEG data was digitally filtered (0.3 Hz high-pass, 50 Hz low-pass) and globally bad channels were detected and removed using FASTER tools (Nolan, Whelan, & Reilly, 2010). To classify a signal as artifactual, FASTER calculates three parameters - variance, mean correlation and Hurst exponent - for each channel; a channel whose data had a Z-score of ±3 for a parameter was deemed to be globally bad. Then, we created a copy of the original dataset, which was then filtered using a 1 Hz high-pass filter and segmented into arbitrary 1 second epochs to identify and remove epochs with excessive artifact. After this, independent component analysis (ICA) was performed on this copied, high-pass filtered dataset; excessive artifacts were detected and removed if amplitude was +/- 1000 uV or if power within the 20-40hz band (after Fourier analysis) was greater than 30dB. After performing an ICA decomposition, ICA weights were copied back to the original dataset; components associated with ocular or other artifacts were identified and removed. Data were epoched to the response markers from −1000 to 2000 ms and baseline corrected using the −400 to −200 ms period preceding the response. Epochs with residual ocular artifact were identified and removed using a +/- 125 μV threshold based on the ocular channels. For all remaining channels, for epochs for which a given channel exhibited voltage +/-125 μV, data were removed and interpolated using a spherical spine interpolation, unless more than 10% of channels were bad within a given epoch, in which case the entire epoch was removed. Finally, any channels that were marked as bad throughout the entire recording were interpolated (spherical spline interpolation) and an average reference was computed. Participants with less than 50% accuracy for Go trials were excluded from analysis (n=3 who were not included in our final sample of 35 children). Similarly, any additional participants with fewer than 4 error trials were excluded from all analyses, similar to criteria used in other studies with the same task for preschool-aged children (e.g., Steele et al., 2016); out of an original sample of 43 children, 8 were not included in our final sample of 35 children. An average of 27.11 error and 138.06 correct trials were available for analysis for participants; an iterative subsampling procedure was employed in order to match the effective trial counts across conditions and participants.

### ERP/EEG measures of inhibitory control and response monitoring

#### ERN/Pe

For ERP analyses, data were down-sampled to 128 Hz to improve processing speed and conserve storage space without any loss of signal quality. ERN was quantified as mean amplitude at a cluster of frontocentral electrodes (Electrodes 7, 4, 54; Figure 2) during an approximately 100 ms window (14 samples; 109.38 ms) immediately following the response; Pe was quantified as mean amplitude at a cluster of centroparietal electrodes (Electrodes 34, 33, 36, 38) during an approximately 200-500 ms post-response window (39 samples; 304.69 ms). These analysis locations and time windows are based on pilot work (S.H. Kim et al., 2017) and previous literature (Grammer et al., 2014; M. Kim et al., 2016); exact time windows reflect actual time resolution possible at the sampling rate used. Mean amplitudes have been found to be robust to increased background noise and variations in the number of trials (Acton, 2013; Luck & Kappenman, 2011). Further, to account for individual differences in trial counts, we employed a subsampling approach, where the ERP/EEG data from a subset of trials is repeatedly selected (with replacement) and averaged 25 times, before being bootstrapped 100 times to yield estimates of condition-averaged ERPs (see Buzzell et al., 2018). The mean ERN and Pe amplitudes were computed on incorrect-response (No-go) trials, with the difference between amplitudes on error vs. correct trials (ΔERN and ΔPe) calculated by subtracting correct-Go waveforms from error-No-go waveforms (error-No-go minus correct-Go).

**Figure 2.**
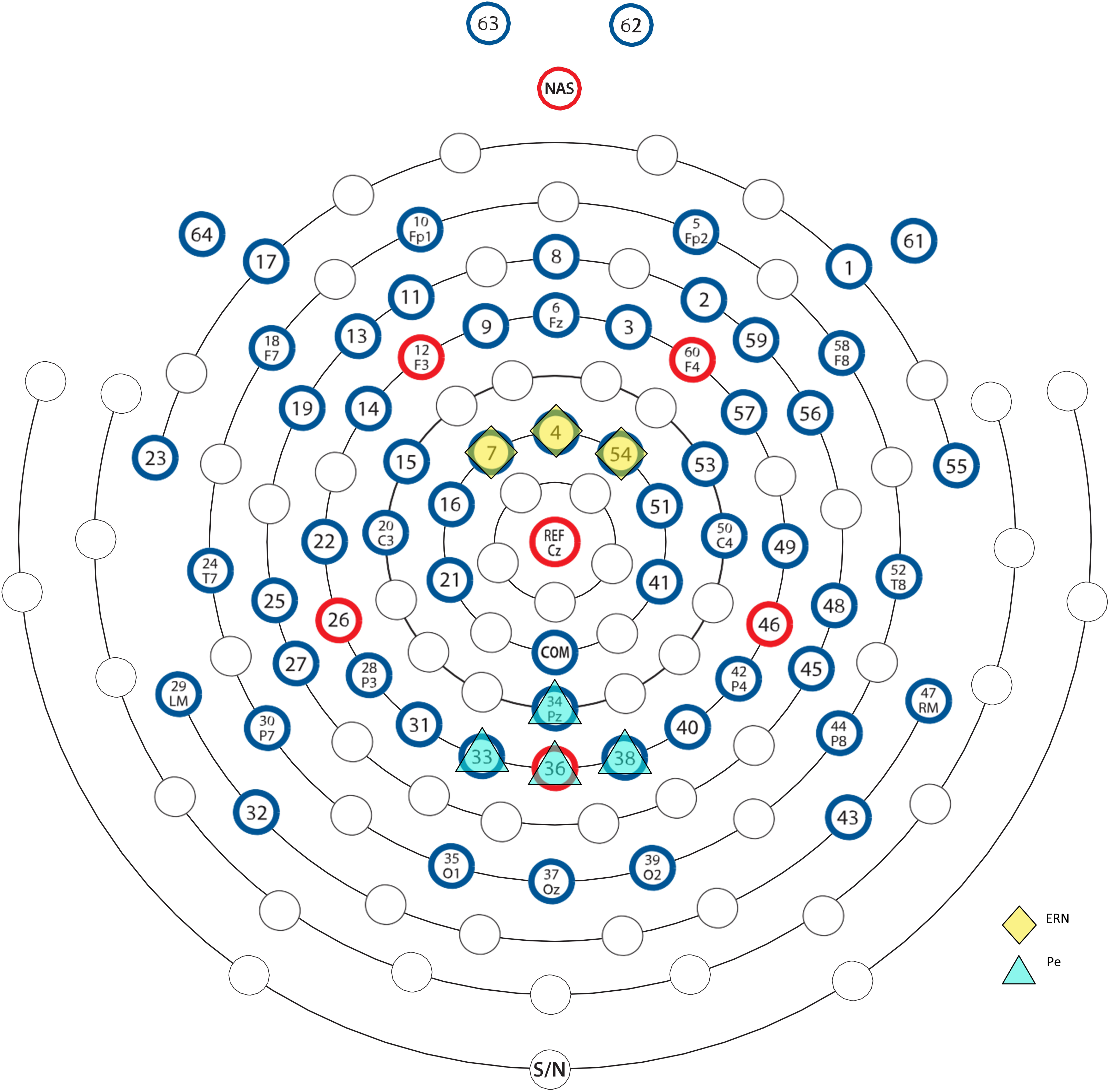
EGI 64-Channel Sensor layout. The EGI 64 electrode HydroCel Geodesic Sensor Net is displayed above. Diamond electrodes indicate fronto-central recording sites for ERN and Theta (Electrodes 7, 4, 54) and triangle electrodes indicate centro-parietal recording sites for Pe (Electrodes 34, 33, 36, 38). The electrode map is used with permission from Electrical Geodesics, Inc. (EGI), Eugene, OR, USA.

#### Frontal error-related theta power

Given our focus on theta (~4-8 Hz) oscillations, we down-sampled the EEG data to 32 Hz in order to improve computational efficiency with no loss of the signals of interest (i.e. Nyquist = 16 Hz). In order to extract error-related theta power, we employed Cohen’s class reduced interference distributions (RIDs) to decompose time frequency (TF) representations of response-locked averaged power (Bernat et al., 2005), after first averaging/bootstrapping trials within each condition of interest to yield equal effective trial counts across conditions. One common approach to decomposing TF representations involves the use of Morlet wavelets (Cohen, 2014); however, the Cohen’s class RID approach produces a time-frequency surface with proven superiority in terms of both time and frequency resolution, as compared to other methods (Bernat et al., 2005). Additionally, in order to isolate error-related theta responses, we performed a principal components analysis (PCA) on the time-frequency surface, focusing on the period from −500 to 500 ms in the 3 hz to 12 Hz time-frequency surface, after first filtering out delta activity; all conditions of interest were entered into the same PCA. Analysis of the scree plot suggested that a 3-factor solution was the best fit to the data. Only one of these factors from the medio-frontal sites reflected a post-response theta component and so further analyses of theta power focused exclusively on this factor. For statistical analyses, factor weights were back projected onto the scalp surface and mean amplitude extracted from a cluster of medio-frontal electrodes (Electrodes 7, 4, 54). The difference between theta power on error vs. correct trials (Δtheta power) was calculated by subtracting correct-Go theta power from error-No-go theta power (error-No-go minus correct-Go). Although the PCA was performed on a time-frequency surface that spanned 3 to 12 Hz and −500 to 500 ms, the actual theta factor that was identified and employed in subsequent analyses was much more focal in both time and frequency, resembling a ~4-8 Hz band effect following the response. The PCA was run on a wider, 3-12 Hz range because the exact upper/lower boundaries of the “theta” band can vary across individuals or development. Our PCA approach identified the theta band in a data-driven manner, isolating a response-related cluster of time-frequency data from within the 3-12 Hz surface on which it was run.

### Behavioral Measures

#### Academic achievement

Academic achievement in reading (Letter-Word Identification, Passage Comprehension) and math (Applied Problems, Math Facts Fluency) at the beginning of kindergarten was measured using the Woodcock-Johnson III NU tests of achievement (WJ; Woodcock, McGrew, Mather, & Schrank, 2001). The Letter-Word Identification sub-test measures the word identification skills. Passage Comprehension measures the understanding of written text. Applied Problems measures the ability to analyze and solve math problems. The Math Facts Fluency sub-test measures the ability to solve simple addition, subtraction and multiplication facts quickly. Standard scores (*M*=100, SD=15) from Letter-Word Identification, Passage Comprehension, and Applied Problems were used for analyses. For the Math Facts Fluency domain, more than 50% of children (n=22) did not achieve the basal for which Standard Scores might not be valid; thus, raw scores were used instead in analyses (See Statistical Analyses for more details).

#### Cognitive skills

Cognitive functioning (nonverbal and verbal IQ) was measured by the Differential Ability Scale (DAS; Elliott, 2007). Nonverbal IQ has been found to be more stable in children with ASD than verbal IQ (Bishop, Farmer, & Thurm, 2015). Therefore, NVIQ was used as an estimate of cognitive skills in our statistical models.

### Statistical Analyses

For all behavioral analyses, response time (RT) analyses were restricted to correct trials and data were log-transformed prior to averaging because RT data are known to be positively skewed (Luce, 1986). Raw RT values are reported in the text for ease of interpretation. We also examined proportions of correct and error responses for Go and No-go trials, respectively. To confirm that ERN and theta power at the frontal sites and the Pe at the posterior sites were significantly larger for error vs. correct trials, t-tests were performed (one-sided). With a priori hypotheses based on past literature, we focused our analyses on particular sites (ERN at frontal, Pe at posterior). However, we also present the results from more complete 2 (site: frontal vs. posterior) X 2 (accuracy: error vs. correct) ANOVAs for both the ERN and Pe in the Supplemental Materials; additionally, based on post-hoc inspection of the ERPs, we include in the Supplemental Materials further analyses of the frontal and posterior sites at an earlier time window (−50 to 50 ms), which is consistent with other work studying the ERN and Pe in children (E. Y. Kim, Iwaki, Imashioya, Uno, & Fujita, 2007; M. Kim et al., 2016). The relations between behavioral performance on the task (log RT and accuracy) and medio-frontal ΔERN and posterior ΔPe, as well as frontal Δtheta power were explored through Pearson *r* correlations. False-Discovery Rate (FDR) corrections were applied for multiple comparisons for the t-tests and correlation analyses. Additionally, we performed regression analyses to examine the effects of neural correlates of error monitoring (ΔERN, ΔPe, and Δtheta power) on academic skills based on the WJ Letter-Word Identification, Passage Comprehension, and Applied Problems Standard Scores while controlling for IQ and behavioral performance to examine whether the effects of neural correlates go above and beyond these constructs that are highly associated with academic performance. For the WJ Math Facts Fluency domain for which more than 50% of children were not able to achieve the basal (raw score = 0), we used Tobit regression which is suitable for zero-inflated continuous outcome variables (Long, 1997). The Tobit regression models jointly (a) the probability for the true outcome being less than 0 (and thus truncated to 0) via logistic regression, and (b) the greater than 0 outcomes as a function of the covariates, via regular regression. All analyses were conducted using SPSS Version 24.

## Results

### Behavioral Assessments

As seen in Table 1, our sample of cognitively-able children with ASD showed average Standard Scores for the WJ Letter-Word Identification, Passage Comprehension, and Applied Problems domains (ranging from mean standard scores [m=100, SD=15] of 98-107). As mentioned above, 22 children were not able to achieve the basal on the Math Facts Fluency domain, although a few children showed very high scores; the raw scores for this domain are reported here. For the Zoo Game, overall mean accuracy was 69% (SD=10%), with accuracy for go trials at 75% (SD=11%) and accuracy for nogo trials at 52% (SD=18%). Mean reaction time for correct-go trials was 624 ms (SD=413 ms).

**Table 1.** Behavioral Assessments (N=35)

|  | Mean | SD | Range |
| --- | --- | --- | --- |
| Woodcock Johnson Test of Achievement IV |  |  |  |
| Letter Word Identification Standard Scores | 107.1 | 15.9 | 76 – 123 |
| Passage Comprehension Standard Scores | 108.0 | 15.8 | 66 – 142 |
| Applied Problem Standard Scores | 97.7 | 13.1 | 69 – 120 |
| Math Facts Fluency Raw Scores* | 2.8 | 6.8 | 0 – 27 |
| Zoo Game |  |  |  |
| Overall Accuracy (%) | 69% | 10% | 50 – 85% |
| Accuracy for Go Trials (% of correct responses) | 75% | 11% | 53 – 91% |
| Accuracy for No-Go Trials (% of no responses) | 52% | 18% | 15 – 88% |
| Reaction Time (Correct-go trials; ms) | 624 | 413 | 413 – 811 |

\*Raw scores were used in analyses because more than a half of the children did not achieve basal, for which standard scores are not valid. However, standard scores ranged from 77 to 152 for this group, with mean of 84.3 and SD of 12.1.

### Medio-Frontal ERN

Figure 3A shows a negative deflection around the time of error commission (relative to correct responses) at frontal electrode sites. Average amplitudes for the error and correct trials, as well as the difference between them, can be seen in Table 2. Mean amplitude for error trials at the frontal sites was significantly more negative than for correct trials [*t*(34)= −4.55, *p*=0.001, FDR adjusted *p*=0.002; Table 2]. An additional 2 (site: frontal vs. posterior) X 2 (accuracy: error vs. correct) ANOVA is presented in the Supplemental Materials. Collectively, the pattern of results confirms the presence of an ERN component, consistent with prior literature (Grammer et al., 2014; E. Y. Kim et al., 2007; M. Kim et al., 2016; M. Kim, Marulis, Grammer, Morrison, & Gehring, 2017; S. H. Kim et al., 2017).

**Table 2.** Mean (SD) ERN and Pe amplitudes and theta average power at the fronto-central and central-posterior sites.

| Fronto-central electrode sites | Mean (SD) |
| --- | --- |
| ERN ($\mu\text{V}$) | -1.08 (2.45) |
| CRN ($\mu\text{V}$) | 1.09 (2.18) |
| $\Delta\text{ERN}$ ($\mu\text{V}$) | -2.17 (2.83) |
| PC-Weighted Error Theta Power (a.u.) | 0.05 (0.04) |
| PC-Weighted Correct Theta Power (a.u.) | 0.03 (0.03) |
| PC-Weighted $\Delta$ Theta Power (a.u.) | 0.02 (0.03) |
| Central-posterior electrode sites | Mean (SD) |
| Pe ($\mu\text{V}$) | -0.62 ( 8.43) |
| Pe Correct ($\mu\text{V}$) | -0.17 ( 7.81) |
| $\Delta$ Pe ($\mu\text{V}$) | 9.56 (10.18) |

ERN, error-related negativity; CRN, correct-related negativity; PC, Principal Component; Pe, error positivity. a.u., arbitrary unit (PC-weighted raw power).

**Figure 3.**
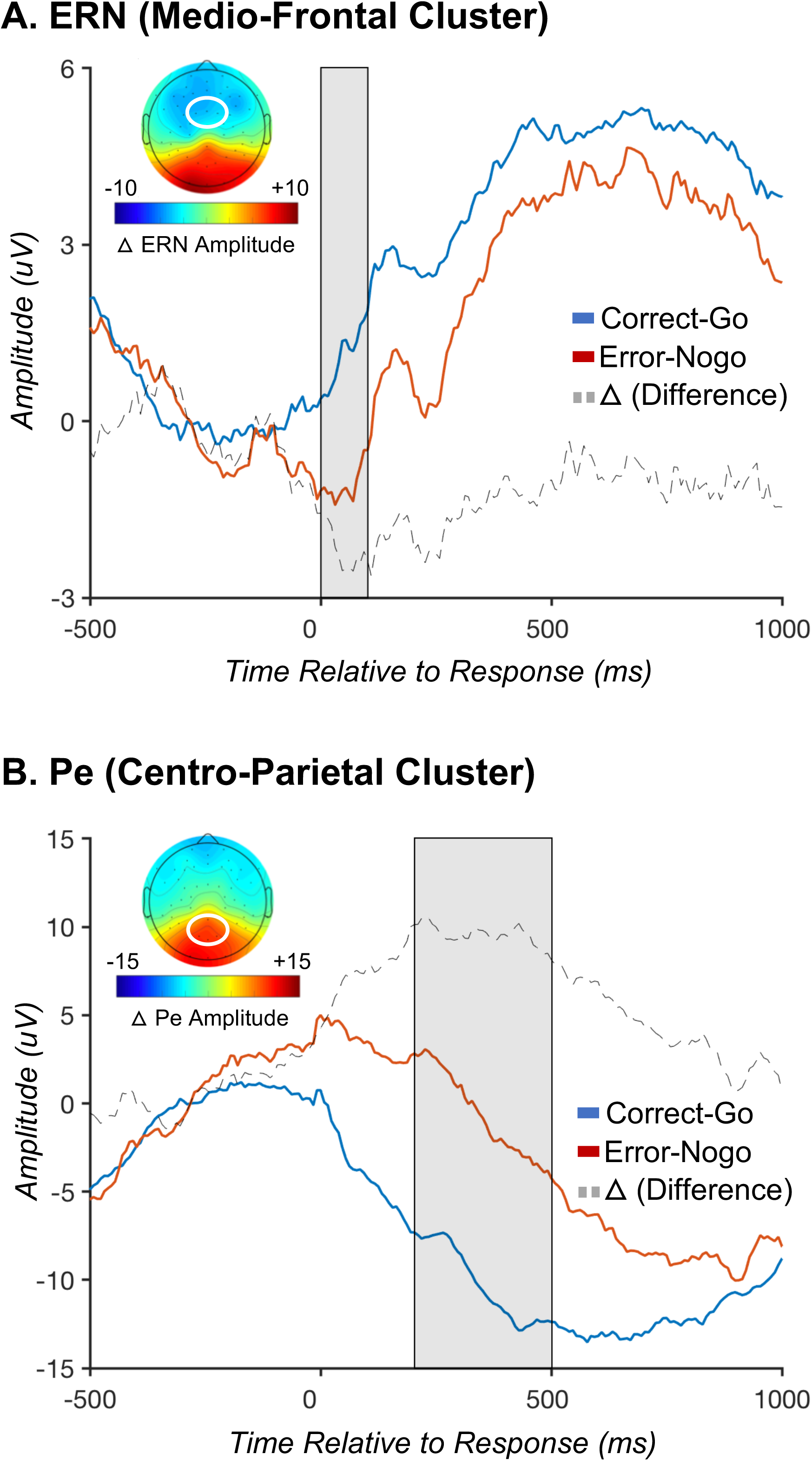
Error-related negativity (ERN) and Error Positivity (Pe). Figure 3A depicts the response-locked ERN at a cluster of frontocentral electrodes, where the ERN is typically maximal. Note that in the ERP plots the ERN is more negative (larger) on error trials (red) than correct trials (blue); in the topographic plot, the ERN is shown to be most negative over frontal regions, as depicted by in blue shading of the topo plot in this region (corresponding to negative values). Dotted line is for ΔERN. Figure 3B depicts the response-locked Pe at a cluster of centroparietal electrodes, where the Pe is typically maximal. Note that the Pe is more positive (larger) on error trials.

### Posterior Pe

Examination of Figure 3B revealed the presence of the Pe at central-posterior electrode sites. Mean amplitude for error trials at the posterior sites was significantly more positive than for correct trials [t(34)= −4.55, p=0.001, FDR adjusted *p*=0.002; Table 2]. An additional 2 (site: frontal vs. posterior) X 2 (accuracy: error vs. correct) ANOVA is presented in the Supplemental Materials. Collectively, the pattern of results confirm the presence of a Pe component, consistent with prior literature (Grammer et al., 2014; E. Y. Kim et al., 2007; M. Kim et al., 2016; M. Kim et al., 2017; S. H. Kim et al., 2017).

### Time Frequency PC-Weighted Theta Power

Figure 4 demonstrates increased theta power (4-8 Hz) on error trials compared to correct trials at the medio-frontal electrode sites. Based on the mean and SD of average theta power, two outlier cases (+/-2.5 SDs) were excluded from all analyses to minimize the effects of those scores on the correlation and regression analyses; these participants were only deemed as outliers for the analyses of theta, but not for the ERPs or behavior, as outlier detection and removal was performed separately for each dependent variable of interest. A t-test showed that theta average power was significantly larger for error trials (compared to correct trials) at the medio-frontal sites [t(1,33=-2.351, p=0.03, FDR adjusted *p*=0.03; Table 2]. This pattern of results for error-related theta is consistent with prior work in adults (Cavanagh & Frank, 2014).

**Figure 4.**
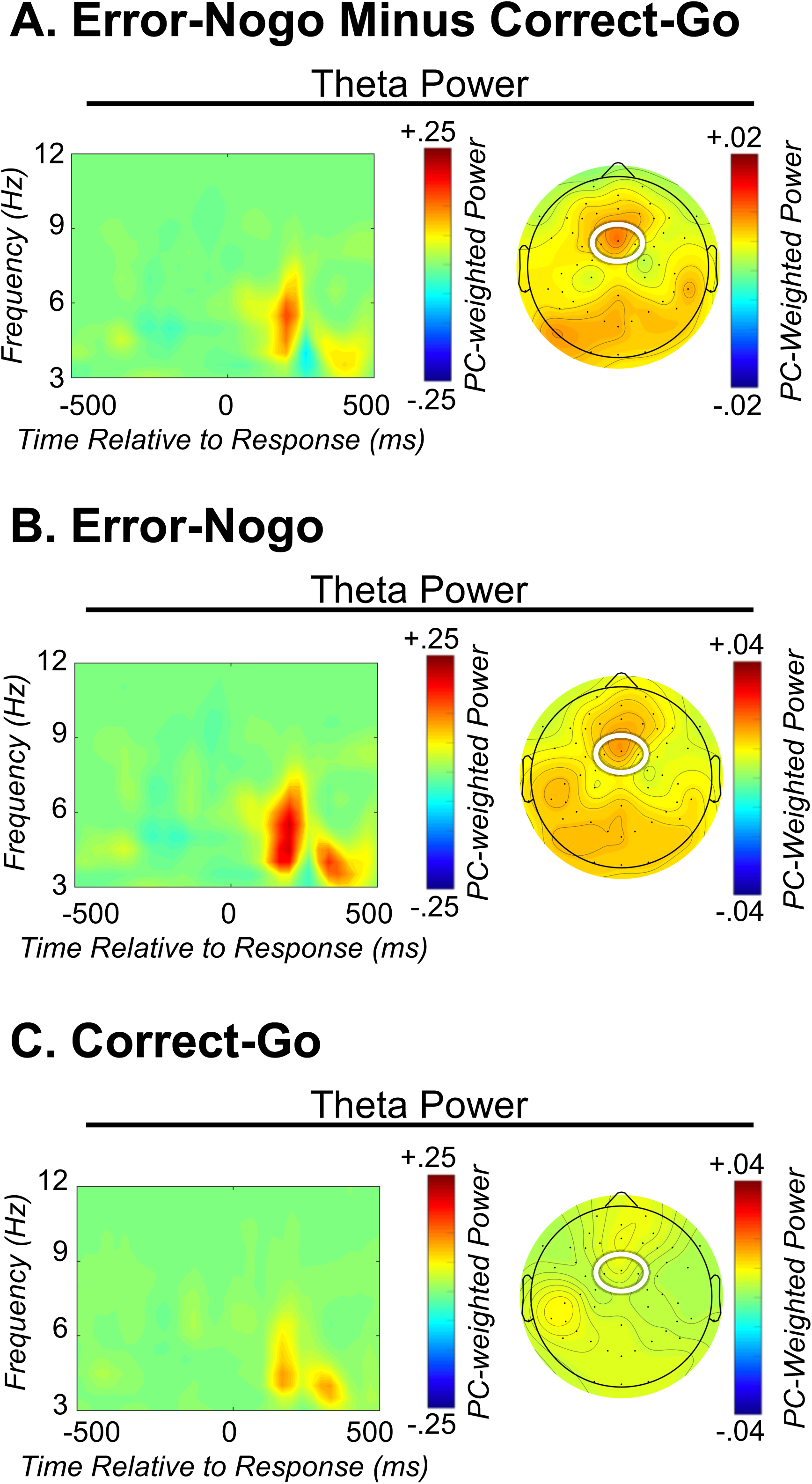
Error-related time-frequency theta power. Figure 4A depicts response-locked theta power after subtraction of correct from error trials and weighting by a principal component that captures error-related theta power. Figure 4B and 4C depict response-locked theta power during error and correct trials respectively. Note that theta power is increased on error trials. Left plots: time-frequency surface plotted for a cluster of frontocentral electrodes. Right plots: corresponding topographic plot of the principal component capturing theta power.

### Associations between ERP/EEG Measures and Behavioral Performance on the Zoo Game

As expected, a more negative frontal ΔERN, a larger posterior ΔPe, and increased frontal Δtheta average power were all significantly correlated with higher overall accuracy (although the association with ΔERN and overall accuracy did not survive the FDR correction; Table 3). Larger frontal Δtheta average power was also significantly correlated with higher go accuracy. Larger posterior ΔPe was significantly correlated with longer reaction times overall and correct-go trials in particular.

**Table 3.** Correlations between ERP/EEG measures and behavioral performance on the Zoo Game

| | Frontal $\Delta ERN$<br>(n=35) | Posterior $\Delta Pe$<br>(n=35) | Frontal $\Delta Theta$<br>Average Power<br>(n=33) |
| --- | --- | --- | --- |
| Overall accuracy | -0.35*(n.s.) | 0.61** | 0.52**(*) |
| Go accuracy | -0.21 | 0.22 | 0.44* |
| No-go accuracy | -0.34* | 0.67** | 0.27 |
| Overall Reaction Time (log transformed) | -0.26 | 0.53** | 0.01 |
| Correct-go Reaction Time (log transformed) | -0.31 | 0.56** | 0.02 |
| Nonverbal IQ | -0.35* | 0.15 | 0.03 |

\* $p < .05$ , \*\* $p < .01$ ; (FDR Adjusted $p$ values reported when different from unadjusted $p$ -values)

### ERP/EEG Measures as Predictors of Academic Skills

As seen in Table 4, regression analyses showed that after controlling for overall accuracy on the Zoo Game and NVIQ, increased Δtheta average power significantly predicted better reading (WJ Passage Comprehension) and math (WJ Math Facts Fluency). *R^2^* significantly improved from 0.19 to 0.3 when Δtheta average power was included in the models to predict Passage Comprehension (p<0.05). For Math Facts Fluency, chi-square analysis of the log-likelihood was performed to confirm that the addition of Δtheta average power significantly improved the prediction of the model (p<0.001). ΔPe also significantly predicted Math Facts Fluency while controlling for accuracy and IQ, and the addition of ΔPe significantly improved the prediction of the model based on the chi-square analysis of the log-likelihood (p<0.05). ERN did not significantly predict any of the academic domains.

**Table 4.**
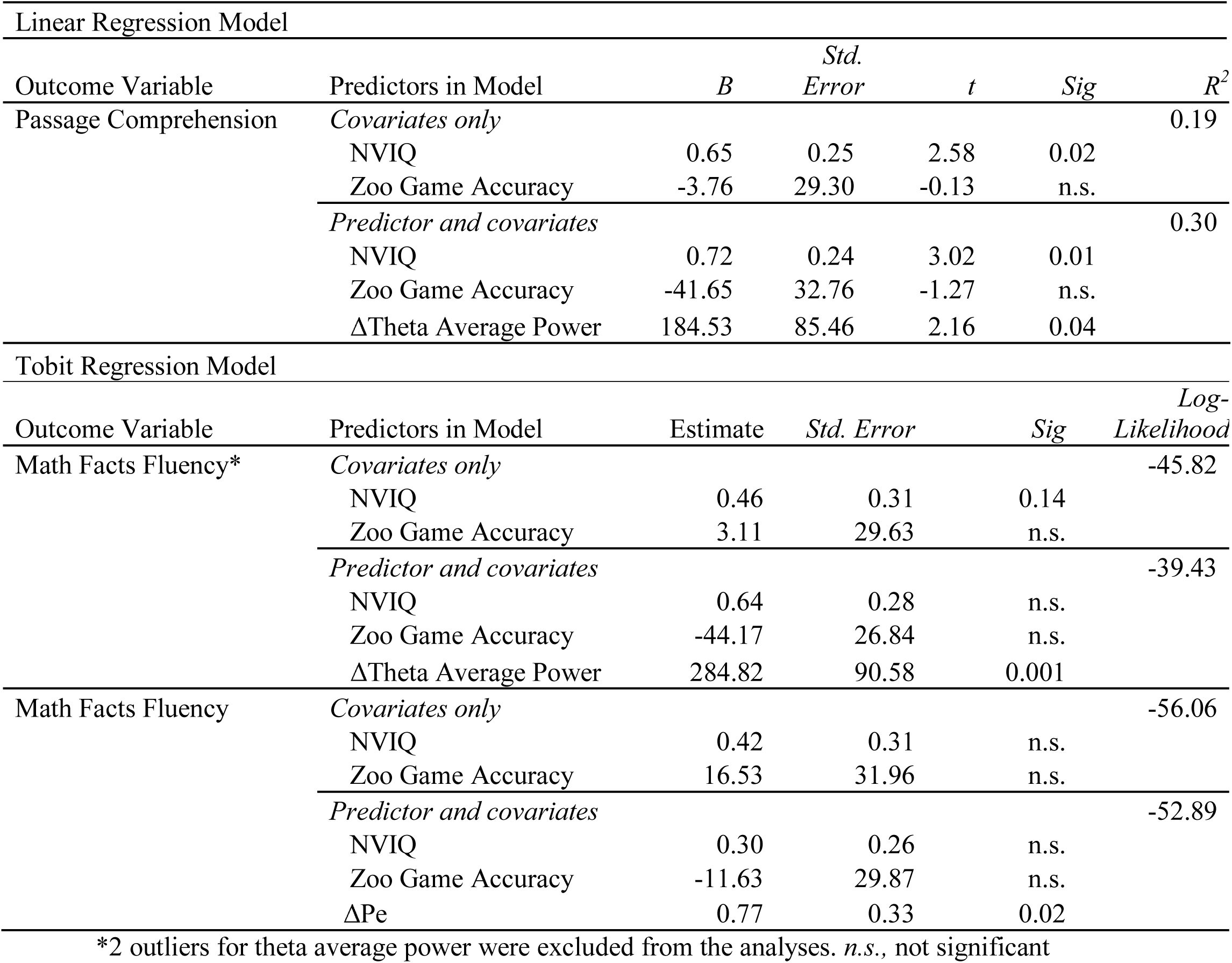
Regression of EEG/ERP measures as predictors of academic skills

## Discussion

Studies based on behavioral assessments of EF have shown that a broad range of EF skills play a key role in the development of academic ability in TD children as well as in children with ASD (Allan, Hume, Allan, Farrington, & Lonigan, 2014; Blair, 2002, 2016; Cameron et al., 2012; Fuster, 1997; Happé et al., 2006; McClelland et al., 2006, 2007; Miyake et al., 2000; Morrison, Ponitz, & McClelland, 2010; Passolunghi & Costa, 2016; Ponitz et al., 2009; Raver, Smith-Donald, Hayes, & Jones, 2005; Wiebe, Espy, & Charak, 2008; Willoughby et al., 2012). Using traditional (ERP) and advanced (time-frequency) electrophysiological methods, we extracted neural activity related to error monitoring—a specific construct falling within EF reflective of one’s ability to detect and process errors while engaging in cognitive tasks—in cognitively-able kindergarteners with ASD. Using the child friendly Go/No-go Zoo Game, we were able to observe the ERN and increased theta power for error trials at medio-frontal sites and Pe at posterior sites in these children. Moreover, we found that a larger Pe and increased theta power were both associated with concurrent academic skills in these young children with ASD at school entry.

Our findings on the presence of ERN and Pe in children with ASD as young as 5 years during a child-friendly Go/No-go task are consistent with past studies based on typically developing children as young as 2-3 years (Abundis-Gutiérrez, Checa, Castellanos, & Rosario Rueda, 2014; Barry & De Blasio, 2015; Ciesielski, Harris, & Cofer, 2004; Grammer et al., 2014) and in older school-aged children with ASD (Henderson et al., 2006; Kemner, Verbaten, Cuperus, Camfferman, & Van Engeland, 1994; Santesso et al., 2011; Sokhadze et al., 2012; South, Larson, Krauskopf, & Clawson, 2010; Vlamings, Jonkman, Hoeksma, van Engeland, & Kemner, 2008). To our knowledge, our study is one of the first to observe ERN and Pe in young children with ASD under 8 years and its connection to behavioral performance (accuracy rate) on the ERP task, building on our prior work with a smaller, independent sample (S.H. Kim et al., 2017).

This study is also the first to successfully employ time-frequency measures to study error monitoring in young children with ASD and to detect significantly increased medio-frontal theta power following error (compared to correct) responses. These data from our children with ASD are consistent with prior work in TD adults and adolescents suggesting that increased theta power over medio-frontal cortex (MFC) underlies error monitoring (Cavanagh & Frank, 2014). It is also worth noting that increased theta power during error trials in the cognitively-able kindergarteners with ASD was significantly correlated with higher levels of task performance (accuracy rates) on the Go/No-go task. However, it remains unclear if increased accuracy rates (and a lower frequency of errors) are what drive increased theta power. In adults, medio-frontal theta power has been shown to arise, at least in part, from ACC, a cortical region that is not only sensitive to errors and conflict, but also to infrequent events (Brown & Braver, 2005; Wessel & Aron, 2017). Thus, the correlation between accuracy rates and theta power in this sample of young children with ASD is consistent with a broader literature on theta power and medio-frontal cortex more generally. The current results provide the first evidence of increased error-related theta power and the expected associations with error frequency, in a sample of children with ASD as young as 5 years.

As hypothesized, we found a significant association between Pe (but not ERN) and math achievement even when NVIQ and the Go/No-Go task accuracy were controlled for, consistent with another study with TD preschoolers (M. Kim et al., 2016). In addition, we found that increased theta power, which may reflect a more domain-general response to errors, was related to both reading and math skills. In contrast, the Pe component has been suggested to reflect more deliberative and task-specific aspects of error processing (Overbeek et al., 2005; Steinhauser & Yeung, 2010), and increased Pe was specifically associated with math skills, but not reading. In addition, a past study in TD preschoolers (M. Kim et al., 2016) found that increased Pe significantly predicted math skills for children performing at or near grade level, but not for those performing well above grade level. Critically, in our current sample of children with ASD, Math Facts Fluency was the domain that showed the lowest average score, with most children scoring in the below average to average range. In turn, it was the only domain that related to the Pe in our sample. These results suggest that the association between the neural correlates of error monitoring and academic outcomes may vary by the level of the child’s functioning as well as the domains of academic skills. Nevertheless, the results support the need to examine EF-related neural activity using complementary approaches, including ERPs and more direct measures based on the time-frequency approach, to predict additional variance in academic achievement for children with ASD, beyond behavioral performance on the Go/No-go task and IQ.

These results have implications at both the basic and applied level. First, in our study, use of the child-friendly Go/No-go Zoo Game adapted from a well-validated task developed by Fox, McDermott and colleagues, as well as Grammer and colleagues (Grammer et al., 2014; Lamm et al., 2014; Troller-Renfree, Zeanah, Nelson, & Fox, 2018), enabled us to maximize the child’s ability to be engaged during the EEG/ERP session allowing us to more effectively examine error-related ERPs and time-frequency data in our sample. Second, researchers should note that because time-frequency analyses allow for extraction of power within specific ranges (e.g. ~4-8 Hz for theta), combined with a PCA approach, effects of high or low frequency artifacts are reduced and the signal-to-noise ratio is maximized (Bernat et al., 2005). This approach appears to enhance our ability to observe EF-related neural dynamics that may be missed by traditional ERP approaches that can be highly sensitive to artifacts and noise (Luck & Kappenman, 2011). Thus, the use of time-frequency analyses, particularly when paired with a PCA approach, may provide new opportunities to investigate neurobiological mechanisms of EF-related cognitive processes more effectively and accurately, especially in young children and those with special needs. Finally, past studies have shown that behavioral performance on EF tasks, including working memory and inhibitory control tasks, is significantly more impaired in children with ASD than in TD peers (Happé et al., 2006; Konstantareas & Stewart, 2006). This finding has prompted the development of targeted interventions aimed at improving EF skills in school-age children (e.g., Kenworthy et al., 2014). Similarly, the results from the present study on the significant link between EF-related neural activity and concurrent academic outcomes in kindergarteners with ASD, combined with our previous work showing atypical EF-related neural activity in young children with ASD compared to matched typical controls (S.H. Kim et al., 2017), point to the importance of incorporating strategies to improve error and conflict monitoring in early autism interventions to maximize academic outcomes.

### Limitations and future direction

Our sample was focused on cognitively-able children with ASD with average to above average cognitive skills without notable structural language delays. This limits the generalizations of our results to children with significant language and cognitive delays. Moreover, math skills in this sample of children showed a wide range of variability with more than the half of the sample not achieving basal scores and a few children showing exceptional computational skills. We were able to find a statistical solution to address the issue of a zero-inflated outcome variable; however, replications with larger, independent, and more representative samples with other developmentally appropriate measures will be important. Furthermore, studies relating the EF-related EEG/ERP measures, specifically those employing time-frequency methods, are still limited in young children, even for typically developing samples. Therefore, a direct comparison between children with ASD and TD children matched on factors such as IQ, performance level on ERP/EEG tasks, and gender, will provide further insights into the examination of atypical neural activity related to error monitoring, or more broadly, to EF.

It will also be important for future research to examine the effects of classroom placements on the development of EF and academic functioning in children with ASD. This might require careful consideration of statistical models and research design because classroom placements are not random but confounded by other factors (e.g., children with lower IQ and language levels being placed in special education classrooms vs. more cognitively able-children in general education or inclusion classrooms). Moreover, even though in our study every effort was made to minimize the artifacts and noise during the EEG/ERP data collection, we included children whose accuracy level for go trials was as low as 50% in an effort to not create a biased sample through over-exclusion based on performance, although this was consistent with past ERP studies with individuals with ASD (e.g., Uzefovsky, Allison, Smith, & Baron-Cohen, 2016). Because the Zoo Task provides children with general feedback after each block to keep them engaged and motivated, future research should also explore potential differences in trial-by-trial feedback-related behavioral performance and neural activity between children with ASD and TD children. We also acknowledge that it is difficult to fully dissociate error-related effects from NoGo-related inhibitory processes when using a Go/NoGo task to study error monitoring, although by selecting incorrect NoGo trials, we chose trials without inhibition (correct Go/incorrect No-Go both involve a motor response and do not involve inhibition, or at least successful inhibition). Nevertheless, despite this limitation, we employed a child-friendly, engaging Go/No-Go task, since Go/No-Go tasks have been widely validated in developmental research examining cognitive control across the life span (Williams, Ponesse, Schachar, Logan, & Tannock, 1999) and well tolerated even by young children. Finally, our focus was to examine the link between the EEG/ERP measures of error monitoring and concurrent academic achievement, however, it will be important to investigate the degree to which EEG/ERP measures *prospectively* predict future academic development. For these reasons, replication with a larger sample of children with ASD as well as matched TD controls using a longitudinal design will be important.

### Conclusion

Our results suggest that the use of time-frequency EEG analyses, complemented by traditional ERP measures, may provide new opportunities to investigate neurobiological mechanisms of EF underlying achievement in young children with ASD. Using advanced time-frequency methods, we found increased response-related theta power during error relative to correct trials in children with ASD as young as 5 years, in addition to traditional response monitoring ERPs, ERN and Pe. More importantly, increased theta power and Pe predicted concurrent academic skills even when controlling for behavioral performance on the EF task and IQ. Results highlight the need to target EF even prior to school entry to maximize academic outcomes in children with ASD.

## Acknowledgements

We thank the children and the families who participated in the study. We also thank the research colleagues who supported the project and this paper: Nurit Benrey, B.A., Shanping Qiu, M.A., Bethany Vibert, Psy.D., Claire Klein, B.A., and Deidre Pozzuoli, M.A.

## References

Abdul Rahman, A., Carroll, D. J., Espy, K. A., & Wiebe, S. A. (2017). Neural correlates of response inhibition in early childhood: Evidence from a Go/No-Go task. Developmental Neuropsychology, 42(5), 336–350.

Abundis-Gutiérrez, A., Checa, P., Castellanos, C., & Rosario Rueda, M. (2014). Electrophysiological correlates of attention networks in childhood and early adulthood. Neuropsychologia, 57, 78–92.

Acton, A. (2013). Issues in Neuropsychology, Neuropsychiatry, and Psychophysiology: 2013 Edition. ScholarlyEditions.

Agam, Y., Hämäläinen, M. S., Lee, A. K. C., Dyckman, K. A., Friedman, J. S., Isom, M., Manoach, D. S. (2011). Multimodal neuroimaging dissociates hemodynamic and electrophysiological correlates of error processing. Proceedings of the National Academy of Sciences, 108(42), 17556–17561.

Allan, N. P., Hume, L. E., Allan, D. M., Farrington, A. L., & Lonigan, C. J. (2014). Relations between inhibitory control and the development of academic skills in preschool and kindergarten: A meta-analysis. Developmental Psychology, 50(10), 2368.

Barry, R. J., & De Blasio, F. M. (2015). Performance and ERP components in the equiprobable go/no-go task: Inhibition in children. Psychophysiology, 52(9), 1228–1237.

Bernat, E. M., Williams, W. J., & Gehring, W. J. (2005). Decomposing ERP time–frequency energy using PCA. Clinical Neurophysiology, 116(6), 1314–1334.

Bishop, S. L., Farmer, C., & Thurm, A. (2015). Measurement of nonverbal IQ in autism spectrum disorder: Scores in young adulthood compared to early childhood. Journal of Autism and Developmental Disorders, 45(4), 966–974.

Blair, C. (2002). School readiness: Integrating cognition and emotion in a neurobiological conceptualization of children’s functioning at school entry. American Psychologist, 57(2), 111.

Blair, C. (2016). Executive function and early childhood education. Current Opinion in Behavioral Sciences, 10, 102–107.

Blair, C., & Razza, R. P. (2007). Relating effortful control, executive function, and false belief understanding to emerging math and literacy ability in kindergarten. Child Development, 78(2), 647–663.

Botvinick, M. M., Cohen, J. D., & Carter, C. S. (2004). Conflict monitoring and anterior cingulate cortex: An update. Trends in Cognitive Sciences, 8(12), 539–546.

Bowers, M. E., Buzzell, G. A., Bernat, E. M., Fox, N. A., & Barker, T. V. (2018). Time-frequency approaches to investigating changes in feedback processing during childhood and adolescence. Psychophysiology.

Brown, J. W., & Braver, T. S. (2005). Learned predictions of error likelihood in the anterior cingulate cortex. Science, 307(5712), 1118–1121.

Buss, K. A., Dennis, T. A., Brooker, R. J., & Sippel, L. M. (2011). An ERP study of conflict monitoring in 4–8-year old children: Associations with temperament. Developmental Cognitive Neuroscience, 1(2), 131–140.

Buzzell, G. A., Barker, T. V., Troller-Renfree, S. V., Bernat, E. M., Bowers, M. E., Morales, S., Fox, N. A. (2018). Adolescent cognitive control, theta oscillations, and social motivation. BioRxiv, 366831.

Buzzell, G. A., Richards, J. E., White, L. K., Barker, T. V., Pine, D. S., & Fox, N. A. (2017). Development of the error-monitoring system from ages 9–35: Unique insight provided by MRI-constrained source localization of EEG. Neuroimage, 157, 13–26.

Cameron, C. E., Brock, L. L., Murrah, W. M., Bell, L. H., Worzalla, S. L., Grissmer, D., & Morrison, F. J. (2012). Fine motor skills and executive function both contribute to kindergarten achievement. Child Development, 83(4), 1229–1244.

Carter, C. S., Macdonald, A. M., Botvinick, M., Ross, L. L., Stenger, V. A., Noll, D., & Cohen, J. D. (2000). Parsing executive processes: Strategic vs. evaluative functions of the anterior cingulate cortex. Proceedings of the National Academy of Sciences, 97(4), 1944–1948.

Cavanagh, J. F., & Frank, M. J. (2014). Frontal theta as a mechanism for cognitive control. Trends in Cognitive Sciences, 18(8), 414–421.

Ciesielski, K. T., Harris, R. J., & Cofer, L. F. (2004). Posterior brain ERP patterns related to the Go/No-Go task in children. Psychophysiology, 41(6), 882–892.

Cohen, M. X. (2014). Analyzing neural time series data: theory and practice. MIT press.

Delorme, A., & Makeig, S. (2004). EEGLAB: an open source toolbox for analysis of single-trial EEG dynamics including independent component analysis. Journal of Neuroscience Methods, 134(1), 9–21.

Elliott, C. D. (2007). Differential Ability Scales (2nd ed.). San Antonio, TX: Harcourt Assessment.

Falkenstein, M., Hohnsbein, J., Hoormann, J., & Blanke, L. (1991). Effects of crossmodal divided attention on late ERP components. II. Error processing in choice reaction tasks. Electroencephalography and Clinical Neurophysiology, 78(6), 447–455.

Frank, M. J., Woroch, B. S., & Curran, T. (2005). Error-related negativity predicts reinforcement learning and conflict biases. Neuron, 47(4), 495–501.

Fuster, J. M. (1997). The prefrontal cortex: Anatomy, physiology, and neuropsychology of the frontal lobe. Lippincott-Raven.

Gehring, W. J., Goss, B., Coles, M. G., Meyer, D. E., & Donchin, E. (1993). A neural system for error detection and compensation. Psychological Science, 4(6), 385–390.

Gotham, K., Pickles, A., & Lord, C. (2009). Standardizing ADOS scores for a measure of severity in autism spectrum disorders. Journal of Autism and Developmental Disorders, 39(5), 693–705.

Grabell, A. S., Olson, S. L., Tardif, T., Thompson, M. C., & Gehring, W. J. (2017). Comparing self-regulation-associated event related potentials in preschool children with and without high levels of disruptive behavior. Journal of Abnormal Child Psychology, 45(6), 1119–1132.

Grammer, J. K., Carrasco, M., Gehring, W. J., & Morrison, F. J. (2014). Age-related changes in error processing in young children: A school-based investigation. Developmental Cognitive Neuroscience, 9, 93–105.

Groen, Y., Wijers, A. A., Mulder, L. J. M., Waggeveld, B., Minderaa, R. B., & Althaus, M. (2008). Error and feedback processing in children with ADHD and children with autistic spectrum disorder: An EEG event-related potential study. Clinical Neurophysiology, 119(11), 2476–2493.

Hajcak, G., McDonald, N., & Simons, R. F. (2003). To err is autonomic: Error-related brain potentials, ANS activity, and post-error compensatory behavior. Psychophysiology, 40(6), 895–903.

Happé, F., Booth, R., Charlton, R., & Hughes, C. (2006). Executive function deficits in autism spectrum disorders and attention-deficit/hyperactivity disorder: Examining profiles across domains and ages. Brain and Cognition, 61(1), 25–39.

Harper, J., Malone, S. M., & Bernat, E. M. (2014). Theta and delta band activity explain N2 and P3 ERP component activity in a go/no-go task. Clinical Neurophysiology, 125(1), 124–132.

Henderson, H., Schwartz, C., Mundy, P., Burnette, C., Sutton, S., Zahka, N., & Pradella, A. (2006). Response monitoring, the error-related negativity, and differences in social behavior in autism. Brain and Cognition, 61(1), 96–109.

Hillman, C. H., Pontifex, M. B., Motl, R. W., O’Leary, K. C., Johnson, C. R., Scudder, M. R., Castelli, D. M. (2012). From ERPs to academics. Developmental Cognitive Neuroscience, 2, S90–S98.

Hirsh, J. B., & Inzlicht, M. (2010). Error-related negativity predicts academic performance. Psychophysiology, 47(1), 192–196.

Jonkman, L. M. (2006). The development of preparation, conflict monitoring and inhibition from early childhood to young adulthood; a Go/Nogo ERP study. Brain Research, 1097(1), 181–193.

Kasari, C., Gulsrud, A., Paparella, T., Hellemann, G., & Berry, K. (2015). Randomized comparative efficacy study of parent-mediated interventions for toddlers with autism. Journal of Consulting and Clinical Psychology, 83(3), 554–563.

Kemner, C., Verbaten, M. N., Cuperus, J. M., Camfferman, G., & Van Engeland, H. (1994). Visual and somatosensory event-related brain potentials in autistic children and three different control groups. Electroencephalography and Clinical Neurophysiology/Evoked Potentials Section, 92(3), 225–237.

Kena, G., Musu-Gillette, L., Robinson, J., Wang, X., Rathbun, A., Zhang, J.,Velez, E. D. V. (2015). The Condition of Education 2015. NCES 2015-144. National Center for Education Statistics.

Kenworthy, L., Anthony, L. G., Naiman, D. Q., Cannon, L., Wills, M. C., Luong-Tran, C., Bal, E. (2014). Randomized controlled effectiveness trial of executive function intervention for children on the autism spectrum. Journal of Child Psychology and Psychiatry, 55(4), 374–383.

Kim, E. Y., Iwaki, N., Imashioya, H., Uno, H., & Fujita, T. (2007). Error-related negativity in a visual go/no-go task: children vs. adults. Developmental Neuropsychology, 31(2), 181–191.

Kim, M., Grammer, J., Marulis, L., Carrasco, M., Morrison, F., & Gehring, W. (2016). Early math and reading achievement are associated with the error positivity. Developmental Cognitive Neuroscience, 22, 18–26.

Kim, M. H., Marulis, L. M., Grammer, J. K., Morrison, F. J., & Gehring, W. J. (2017). Motivational processes from expectancy–value theory are associated with variability in the error positivity in young children. Journal of Experimental Child Psychology, 155, 32–47.

Kim, S. H., Bal, V. H., & Lord, C. (2017). Longitudinal follow-up of academic achievement in children with autism from age 2 to 18. Journal of Child Psychology and Psychiatry.

Kim, S. H., Grammer, J., Benrey, N., Morrison, F., & Lord, C. (2018). Stimulus processing and error monitoring in more-able kindergarteners with autism spectrum disorder: A short review and a preliminary Event-Related Potentials study. European Journal of Neuroscience, 47(6), 556–567.

Konstantareas, M. M., & Stewart, K. (2006). Affect regulation and temperament in children with autism spectrum disorder. Journal of Autism and Developmental Disorders, 36(2), 143–154.

Kramer, J. H., Mungas, D., Possin, K. L., Rankin, K. P., Boxer, A. L., Rosen, H. J., Widmeyer, M. (2014). NIH examiner: Conceptualization and development of an executive function battery. Journal of the International Neuropsychological Society□: JINS, 20(1), 11–19.

Lamm, C., Walker, O. L., Degnan, K. A., Henderson, H. A., Pine, D. S., McDermott, J. M., & Fox, N. A. (2014). Cognitive control moderates early childhood temperament in predicting social behavior in 7-year-old children: an ERP study. Developmental Science, 17(5), 667–681.

Larson, M. J., Clayson, P. E., & Clawson, A. (2014). Making sense of all the conflict: A theoretical review and critique of conflict-related ERPs. International Journal of Psychophysiology, 93(3), 283–297.

Long, J. S. (1997). Regression models for categorical and limited dependent variables (Vol. 7). Thousand Oaks: Sage Publications.

Lord, C., Rutter, M., DiLavore, P. C., Risi, S., Gotham, K., & Bishop, S. (2012). Autism Diagnostic Observation Schedule: ADOS-2. Los Angeles, CA: Western Psychological Services.

Luce, R. D. (1986). Response times: Their role in inferring elementary mental organization. Oxford University Press on Demand.

Luck, S. J., & Kappenman, E. S. (2011). The Oxford Handbook of Event-Related Potential Components. Oxford University Press.

McClelland, M. M., Acock, A. C., & Morrison, F. J. (2006). The impact of kindergarten learning-related skills on academic trajectories at the end of elementary school. Early Childhood Research Quarterly, 21(4), 471–490.

McClelland, M. M., Cameron, C. E., Connor, C. M., Farris, C. L., Jewkes, A. M., & Morrison, F. J. (2007). Links between behavioral regulation and preschoolers’ literacy, vocabulary, and math skills. Developmental Psychology, 43(4), 947.

Miyake, A., Friedman, N. P., Emerson, M. J., Witzki, A. H., Howerter, A., & Wager, T. D. (2000). The unity and diversity of executive functions and their contributions to complex “Frontal Lobe” tasks: A latent variable analysis. Cognitive Psychology, 41(1), 49–100.

Morrison, F. J., Ponitz, C. C., & McClelland, M. M. (2010). Self-regulation and academic achievement in the transition to school. In Child development at the intersection of emotion and cognition (pp. 203–224). Washington, DC, US: American Psychological Association.

Nolan, H., Whelan, R., & Reilly, R. B. (2010). FASTER: Fully automated statistical thresholding for EEG artifact rejection. Journal of Neuroscience Methods, 192(1), 152–162.

Overbeek, T. J., Nieuwenhuis, S., & Ridderinkhof, K. R. (2005). Dissociable components of error processing: On the functional significance of the Pe vis-à-vis the ERN/Ne. Journal of Psychophysiology, 19(4), 319–329.

Passolunghi, M. C., & Costa, H. M. (2016). Working memory and early numeracy training in preschool children. Child Neuropsychology, 22(1), 81–98.

Ponitz, C. C., McClelland, M. M., Matthews, J. S., & Morrison, F. J. (2009). A structured observation of behavioral self-regulation and its contribution to kindergarten outcomes. Developmental Psychology, 45(3), 605–619.

Posner, M. I., & Raichle, M. E. (1994). Images of mind. Scientific American Library/Scientific American Books.

Raver, C. C., Smith-Donald, R., Hayes, T., & Jones, S. M. (2005). Self-regulation across differing risk and sociocultural contexts: Preliminary findings from the Chicago School Readiness Project. In biennial meeting of the Society for Research in Child Development, Atlanta, GA.

Rogers, S., & Dawson, G. (2010). Early start Denver model for young children with autism: Promoting language, learning, and engagement. Guilford Press.

Santesso, D. L., Drmic, I. E., Jetha, M. K., Bryson, S. E., Goldberg, J. O., Hall, G. B., Schmidt, L. A. (2011). An event-related source localization study of response monitoring and social impairments in autism spectrum disorder. Psychophysiology, 48(2), 241–251.

Schneider, W. (2010). Metacognition and memory development in childhood and adolescence. Metacognition, Strategy Use, and Instruction, 54–81.

Sokhadze, E. M., Baruth, J. M., Sears, L., Sokhadze, G. E., El-Baz, A. S., & Casanova, M. F. (2012). Prefrontal neuromodulation using rTMS improves error monitoring and correction function in autism. Applied Psychophysiology and Biofeedback, 37(2), 91–102.

South, M., Larson, M. J., Krauskopf, E., & Clawson, A. (2010). Error processing in high-functioning autism spectrum disorders. Biological Psychology, 85(2), 242–251.

Steele, V.R., Anderson, N.E., Claus, E.D., Bernat, E.M., Rao, V., Assaf, M., Pearlson, G.D., Calhoun, V.D., and Kiehl, K.A. (2016). Neuroimaging measures of error-processing: Extracting reliable signals from event-related potentials and functional magnetic resonance imaging. NeuroImage, 132, 247–260.

Steinhauser, M., & Yeung, N. (2010). Decision processes in human performance monitoring. Journal of Neuroscience, 30(46), 15643–15653.

Taylor, S. F., Stern, E. R., & Gehring, W. J. (2007). Neural systems for error monitoring recent findings and theoretical perspectives. The Neuroscientist, 13(2), 160–172.

Troller-Renfree, S., Zeanah, C. H., Nelson, C. A., & Fox, N. A. (2018). Neural and cognitive factors influencing the emergence of psychopathology: Insights from the Bucharest early intervention project. Child Development Perspectives, 12(1), 28–33.

Ullsperger, M., Fischer, A. G., Nigbur, R., & Endrass, T. (2014). Neural mechanisms and temporal dynamics of performance monitoring. Trends in Cognitive Sciences, 18(5), 259–267.

Uzefovsky, F., Allison, C., Smith, P., & Baron-Cohen, S. (2016). Brief report: The Go/No-Go task online: Inhibitory control deficits in autism in a large sample. Journal of Autism and Developmental Disorders, 46(8), 2774–2779.

Vlamings, P. H. J. M., Jonkman, L. M., Hoeksma, M. R., van Engeland, H., & Kemner, C. (2008). Reduced error monitoring in children with autism spectrum disorder: An ERP study. European Journal of Neuroscience, 28(2), 399–406.

Wessel, J. R., & Aron, A. R. (2017). On the globality of motor suppression: Unexpected events and their influence on behavior and cognition. Neuron, 93(2), 259–280.

Wetherby, A. M., Guthrie, W., Woods, J., Schatschneider, C., Holland, R. D., Morgan, L., & Lord, C. (2014). Parent-implemented social intervention for toddlers with autism: An RCT. Pediatrics, 134(6), 1084–1093.

Wiebe, S. A., Espy, K. A., & Charak, D. (2008). Using confirmatory factor analysis to understand executive control in preschool children: I. Latent structure. Developmental Psychology, 44(2), 575.

Williams, B. R., Ponesse, J. S., Schachar, R. J., Logan, G. D., & Tannock, R. (1999). Development of inhibitory control across the life span. Developmental Psychology, 35(1), 205.

Willoughby, M. T., Kupersmidt, J. B., & Voegler-Lee, M. E. (2012). Is preschool executive function causally related to academic achievement? Child Neuropsychology, 18(1), 79–91.

Woodcock, R. W., McGrew, K. S., Mather, N., & Schrank, F. (2001). Woodcock-Johnson III: Tests of Achievement. Itasca, IL: Riverside Publishing.

