## Supplement File for "Neural Dynamics of Executive Function in Cognitively-able Kindergarteners with Autism Spectrum Disorders (ASD) as Predictors of Concurrent Academic Achievement"

**Supplemental Material**

**Two (site; frontal vs. posterior) by Two (accuracy: error vs. correct) ANOVA results for ERN and Pe**

In the main text, we specifically analyze the ERN at a frontal scalp location, and the Pe at a posterior scalp location, consistent with prior literature that has established the scalp locations where these ERPs are maximal (Grammer et al., 2014; E. Y. Kim, Iwaki, Imashioya, Uno, & Fujita, 2007; M. Kim et al., 2016; M. H. Kim, Marulis, Grammer, Morrison, & Gehring, 2017; S. H. Kim et al., 2017). However, we also present the results from more complete 2 (site: frontal vs. posterior) X 2 (accuracy: error vs. correct) ANOVAs for both the ERN and Pe here.

**ERN (0-100 ms mean amplitude).** The significant main effect of accuracy (F=13.888, p=0.001) and a significant site and accuracy interaction effect (F=27.394, p<0.001) emerged. For the frontal sites, as expected, the amplitude for the error trials was more negative than the amplitude for the correct trials (p<0.001), whereas the reverse was found for the posterior sites (p<0.001).

**Pe (200-500 ms mean amplitude).** The significant main effect of site (F=35.358, p<0.001) and accuracy (F=31.510, p<0.001), and a significant site and accuracy interaction effect (F=26.875, p<0.001) emerged. The difference in the amplitudes between correct and error trials in the posterior sites was significantly larger (and more positive) than the difference in the frontal sites (p<0.001).

**ERP analyses based on -50 to 50 ms window**

In analyzing the ERN, we chose to utilize a conservative 0 – 100 ms window, based on a larger body of literature studying the ERN (Grammer et al., 2014; E. Y. Kim, Iwaki, Imashioya, Uno, & Fujita, 2007; M. Kim et al., 2016; M. H. Kim, Marulis, Grammer, Morrison, & Gehring, 2017; S. H. Kim et al., 2017); similarly, based on prior work we analyzed, the Pe using a 200 – 500 ms window (Grammer et al., 2014; E. Y. Kim et al., 2007; M. Kim et al., 2016; M. H. Kim et al., 2017; S. H. Kim et al., 2017). Nonetheless, post-hoc inspection of the ERPs suggest that the ERN may start prior to 0 ms, and similarly, that the posterior Pe appears to diverge prior to 0 ms. We note here that this pattern of ERP waveforms is quite similar to other work investigating the ERN and Pe in children (Kim, Marulis, Grammer, Morrison, & Gehring, 2017; Kim, Iwaki, Imashioya, Uno, & Fujita, 2007). Therefore, although we retain our a priori analysis time windows and electrode locations for all analyses in the main text, in order to provide a more complete characterization of the ERPs observed in the current study, we further conducted a Two (site; frontal vs. posterior) by Two (accuracy: error vs. correct) ANOVA using the mean amplitude during a -50 to 50 ms window.

*Two (site; frontal vs. posterior) by Two (accuracy: error vs. correct) ANOVA results.* The significant main effects of accuracy (F=6.949, p=0.013) and site (F=6.859, p=0.013) and a significant site and accuracy interaction effect (F=21.042, p<0.001) emerged. For the frontal sites, as expected, the amplitude for the error trials was more negative than the amplitude for the correct trials (p<0.001), whereas the reverse was found for the posterior sites (<0.001). Critically, these analyses are consistent with the ERP results reported in the main text, while also extending the ERP results by demonstrating similarities between the ERPs observed in the current study and other work investigating the ERN and Pe in children at earlier time windows (Kim, Marulis, Grammer, Morrison, & Gehring, 2017; Kim, Iwaki, Imashioya, Uno, & Fujita, 2007); we refer readers to this other work for a more in depth discussion of the ERN and Pe at earlier time windows.
